# Characterization of energy filtering slit widths for MicroED data collection

**DOI:** 10.1101/2025.02.24.639939

**Authors:** Max T.B. Clabbers, Johan Hattne, Michael W. Martynowycz, Tamir Gonen

## Abstract

A favorable signal-to-noise ratio is essential for obtaining high-quality diffraction data in macromolecular electron crystallography. Inelastic scattering contributes significantly to the noise, reducing contrast between diffraction peaks and background, which complicates peak detection and compromises the accuracy of intensity integration. Energy filtering mitigates these challenges and enhances diffraction data quality by removing the inelastically scattered electrons, leading to reduced background noise and sharper Bragg peaks. Previously, we reported a substantial improvement in MicroED data quality and resolution with energy filtering. Here, we systematically evaluate the impact of different energy filter slit widths for optimal MicroED data collection. Data from proteinase K lamellae were collected using the 5, 10, and 20 eV energy filter slit widths. Our results show that the narrowest slit widths result in a stronger diffraction signal with lower background noise, improving the precision of the intensity measurements which resulted in better structural models. Our findings provide insights into the optimization of energy filter slit settings that, when paired with direct electron detection, enhance MicroED data collection strategies in MicroED by improving the signal-to-noise ratio, supporting higher quality data and ultimately enabling more precise structure determination.

## 1. Introduction

Data collection is a critical part of any crystallographic experiment, where a higher signal-to-noise ratio allows for more precise intensity measurements and ultimately leads to better structural models. Optimizing the experimental setup and data collection strategies is therefore essential to maximize the amount of useful structural information that can be extracted from each crystal. In X-ray diffraction, highly accurate data are collected at synchrotron facilities, however radiation damage limits the minimum crystal size that can be used (Sanishvili *et al*., 2011; Holton & Frankel, 2010). In electron crystallography, the signal is generally much weaker because it relies on crystals that are much smaller, resulting in fewer repeating units to amplify the coherent kinematic diffraction signal. Fortunately, electrons deposit substantially less energy into the sample per useful elastic scattering event compared to X-rays (Henderson, 1995). This makes it possible to obtain sufficiently strong and highly accurate data from small crystals in microcrystal electron diffraction (MicroED) (Nannenga *et al*., 2014). Faster and more sensitive detectors allowed continuous rotation data collection, and significantly improved data quality and resolution (Nannenga *et al*., 2014; Yonekura *et al*., 2015; Clabbers *et al*., 2017). More recently, we demonstrated that employing electron counting data collection on direct electron detectors further improves the signal-to-noise ratio (Martynowycz *et al*., 2022; Clabbers *et al*., 2022). Even under low flux density conditions, these cameras are sensitive enough to capture the diffraction signal. Despite these advances, uncertainties in the diffraction geometry and sources of experimental noise persist, suggesting room for further optimization of hardware setups and data collection strategies to achieve higher resolution and improved data quality.

A major source of noise in electron microscopy is inelastic scattering, which can obscure otherwise valuable structural information. Inelastically scattered electrons lose a small fraction of their energy and undergo a phase shift, rendering them incoherent with the primary elastic scattering events (Cowley & Moodie, 1957). Inelastic scattering events therefore interfere with the coherent kinematic diffraction signal that is focused in the Bragg spots. This presents a significant challenge, as inelastic scattering occurs 3-4 times more frequently than elastic scattering in biological specimens (Henderson, 1995; Latychevskaia & Abrahams, 2019). Most inelastically scattered electrons will end up near the direct beam, as the scattering angle for inelastic events is generally relatively small. This leads to high background noise surrounding the beam center, which obscures the low-resolution diffraction spots. This inelastic background noise then rapidly falls off with increasing distance from the center. Additionally, there is a high probability of multiple scattering interactions due to the short inelastic mean-free-path of electrons (Fujiwara, 1959; Latychevskaia & Abrahams, 2019). These multiple scattering events contribute to the background noise, but can also manifest as a broadening of the Bragg peaks. When an elastic event is followed by inelastic scattering the electrons become partially incoherent but they remain focused near the crystal lattice (Latychevskaia & Abrahams, 2019). The increased background and peak broadening both interfere with the data integration, affecting estimation of the intensities of the peak profiles, and complicating the detection of faint high-angle reflections that are at or below the noise level. To make matters worse, these effects can become even more pronounced as the amount of sample the electrons traverse increases, either because of thicker samples or at higher tilt angles.

In electron microscopy, energy filtering can selectively remove inelastic electrons that have lost more than a specific amount of energy, effectively isolating the coherent elastic signal and improving the signal-to-noise ratio. Previous reports assessed the improvements of both in- and post-column energy filters in macromolecular crystallography (Gonen *et al*., 2004, 2005; Yonekura *et al*., 2002, 2015, 2019) and materials science (Gemmi & Oleynikov, 2013; Yang *et al*., 2022). These studies consistently demonstrate a clear reduction in background noise, sharper Bragg peaks, better signal-to-noise ratios, and improved intensity and model statistics. Recently, we demonstrated that energy filtering combined with direct electron detection significantly improves MicroED data quality and resolution compared to unfiltered data (Clabbers *et al*., 2024). To achieve this, we employed a strategy of collecting several small sweeps to accommodate the restraints posed on the flux density per frame of the camera in counting mode and the total fluence that can be withstood by the crystals. This approach improved the resolution of proteinase K lamellae from 1.40 Å for unfiltered data to 1.09 Å resolution for filtered MicroED, resolving detailed structural features including individual hydrogen atoms positions (Clabbers *et al*., 2024). These results underscore the impact of optimizing instrumentation to get the best data quality from the sample, leading to more accurate structural models.

Here, we systemically investigate and compare different energy filter slit width settings to identify the optimal conditions for macromolecular MicroED data collection. Using standardized proteinase K lamellae, we analyzed diffraction stills and MicroED data recorded using the 5, 10, and 20 eV slits to evaluate their effects on the signal-to-noise ratio, intensity measurements, and model statistics.

## 2. Methods

### 2.1. Crystallization

Proteinase K microcrystals were are grown as previously described (Masuda *et al*., 2017; Martynowycz *et al*., 2023; Clabbers *et al*., 2024). Briefly, 40 mg/ml protein solution in 20 mM MES-NaOH pH 6.5 was mixed at a 1:1 ratio with a precipitant solution of 0.5 M NaNO_3_, 0.1 M CaCl_2_, 0.1 M MES-NaOH pH 6.5. The mixture was incubated at 4 °C, and microcrystals with dimensions of approximately 7-12 μm appeared within 24h.

### 2.2. Sample preparation

Holey carbon electron microscopy grids (Quantifoil, Cu 200 mesh, R2/2) were glow discharged for 30 s at 15 mA on the negative setting. Two grids were prepared using a Leica GP2 vitrification device set at 4 °C and 90% humidity. For each sample, 3 μl of crystal solution was deposited onto the grid, incubated for 10s, and any excess liquid was removed from the back side using filter paper. The sample was then soaked on-grid for 30s with 3 μl of a cryoprotectant solution containing 30% glycerol, 250 mM NaNO_3_, 50 mM CaCl_2_, 60 mM MES-NaOH pH 6.5. Any excess liquid was blotted away, and the grid was rapidly vitrified by plunging it in liquid ethane. Grids were stored in a liquid nitrogen dewar.

### 2.3. Focused ion beam milling

Grids were loaded onto a Helios Hydra 5 CX dual-beam plasma FIB/SEM (Thermo Fisher Scientific). The sample was coated with a thin protective layer of platinum for 45s using the gas injection system. Lamellae were made using a 30 kV Argon plasma ion beam following a stepwise protocol as described previously (Martynowycz *et al*., 2023; Clabbers *et al*., 2024). Summarily, coarse milling steps were performed using a 2.0 nA current to a thickness of approximately 3 μm. Finer milling steps at 0.2 nA were used to thin the lamellae to 600 nm. Final polishing steps were performed at 60 pA down to an optimal thickness of 300 nm, equal to approximately one time the inelastic mean free path at 300 kV (Martynowycz *et al*., 2021). In each session, nine lamellae were prepared sequentially using this protocol. Lamellae for the 5 and 10 eV slit setting tests were prepared from the same grid over two sessions. Lamellae for the 20 eV data were prepared on another session from a second grid. After each session, grids were immediately cryo-transferred to the transmission electron microscope (TEM) for data collection. During the transfer step, the grids were rotated by 90° relative to the milling direction such that the rotation axis on the microscope is perpendicular to the milling direction.

### 2.4. Data collection

Diffraction data were collected on a Titan Krios G3i TEM (Thermo Fisher Scientific) equipped with a X-FEG operated at an acceleration voltage of 300 kV, a post-column Selectris energy filter, and a Falcon 4i direct electron detector. The microscope was aligned for low flux density conditions using the 50 μm C2 aperture, spot size 11, gun lens setting 8, and a parallel electron beam of 10 μm diameter. Under these conditions, the flux density was approximately 0.002 e^-^/Å^2^/s. The energy spread of the emitted electrons was characterized as *ΔE* = 0.834 eV +/-0.006 eV at full-width half max (FWHM). The zero-loss peak of the energy filter was aligned in defocused diffraction mode and centered within the SA aperture. Data were collected using the 150 μm selected area (SA) aperture, defining an area with ∼3.5 μm diameter at the sample plane. The detector distance was 1402 mm and calibrated prior to data collection using a standard evaporated aluminum grid (Ted Pella). Diffraction stills were taken of a stationary crystal for brief 10 s exposures at a total fluence of ∼0.02 e^-^ /Å^2^. Data were first collected from the same position without any energy filter slit inserted, followed by sequentially setting the slit width to 5, 10, and 20 eV settings. MicroED data were collected using the continuous rotation method for 420 s exposures at an angular increment of 0.0476 °/s, covering a total rotation range of 20.0° at a fluence of ∼0.84 e^-^/Å^2^. Data were recorded on a Falcon 4i direct electron detector in electron counting mode operating at an internal frame rate of ∼320 Hz. The proactive dose protector was manually disabled. Raw data were written in electron event representation (EER) format with an effective readout speed of ∼308 frames per second, not counting gap frames.

### 2.5. Data processing and structure refinement

Individual MicroED datasets were binned by two and converted to SMV format after applying post-counting gain corrections using the MicroED tools (available at https://cryoem.ucla.edu/downloads). Diffraction stills were summed in batches of 3080 frames, whereas continuous rotation MicroED data were summed in batches of 308, such that each summed image represents a 1 s exposure. Individual MicroED datasets were processed using XDS (Kabsch, 2010). Data were integrated up to a cross-correlation between two random half sets that was still significant at the 0.1% level (Karplus & Diederichs, 2012). Individual datasets were analyzed and merged using XSCALE (Kabsch, 2010). The merged data were truncated at 1.2 Å resolution, enabling a direct comparison between intensity and model statistics for each of the different energy filer slits. Data were merged using Aimless (Evans & Murshudov, 2013) and refined against the same starting model (PDB ID 9DHO) (Clabbers *et al*., 2024). For each slit width setting, structures were refined using the same protocol in phenix.refine (Afonine *et al*., 2012) including electron scattering factors, automated optimization of the geometry and ADP weights, and individual anisotropic *B*-factors for all non-hydrogen atoms.

## 3. Results

### 3.1. Evaluation of energy filter slit widths on the diffraction signal

To evaluate the different energy filter slit settings, diffraction patterns were recorded from the same area of a 300 nm thin proteinase K lamella that was kept stationary in the beam throughout. Each diffraction pattern was collected using a brief 10 s (∼3080 frames) exposure at a total fluence of 0.02 e^-^/Å^2^ by sequentially using no energy filter, followed by inserting the 5, 10, and 20 eV slit settings (**Fig. 1**). Each diffraction still was taken from the same location using a ∼3.5 μm diameter parallel electron beam as defined by the SA aperture. Visual inspection of the diffraction stills shows a substantial reduction in background noise and sharper peaks in all the filtered diffraction patterns compared to the unfiltered data. The better signal-to-noise is especially noticeable for the low-resolution reflections near the center beam position, which become clearly distinguishable from the background when the energy filter is used.

**Figure 1.**
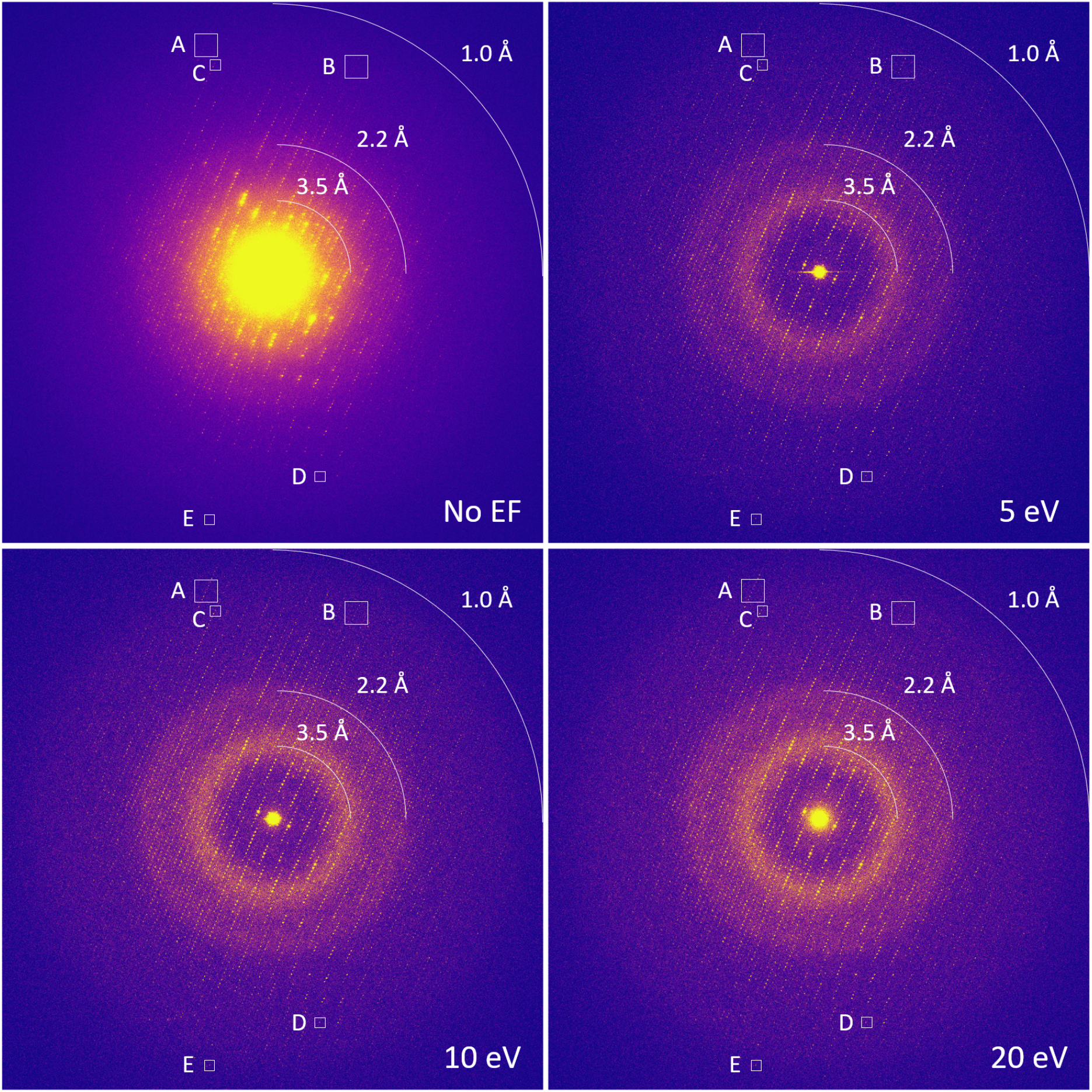
Filtered and unfiltered diffraction patterns of a proteinase K lamella taken especially without the energy filter slit inserted, followed by the 5, 10, and 20 eV slit settings. Each diffraction pattern was taken at 10 s exposure with a total fluence of 0.02 e^-^/Å^2^. Diffraction patterns were taken at the same location on the lamella using a ∼ 3.5 μm parallel beam as defined by the SA aperture. The highlighted areas A-E were used to plot the peak profiles presented in **Figure 2**.

With narrower energy filter slits, the background noise is progressively reduced, enhancing not only the visibility of the lower angle reflections, but making spots at medium and high-resolution clearer, extending the resolution of discernible diffraction to the edge of the detector at ∼1 Å resolution (**Fig. 1**). In comparison, any diffraction spots beyond approximately 1.2 Å resolution are difficult to separate from the background in the unfiltered data. Spot profiles shown at ∼1.2 and ∼1.1 Å resolution further illustrate this improvement, with rows of Bragg peaks appearing clearly above the noise level in unfiltered that were hidden within the much noisier background without energy filtering (**Fig. 2A**). Although filtered diffraction spots are generally less intense than unfiltered data, they nevertheless appear sharper due to the reduction of the background, as illustrated for individual spot profiles (**Fig. 2B**).

**Figure 2.**
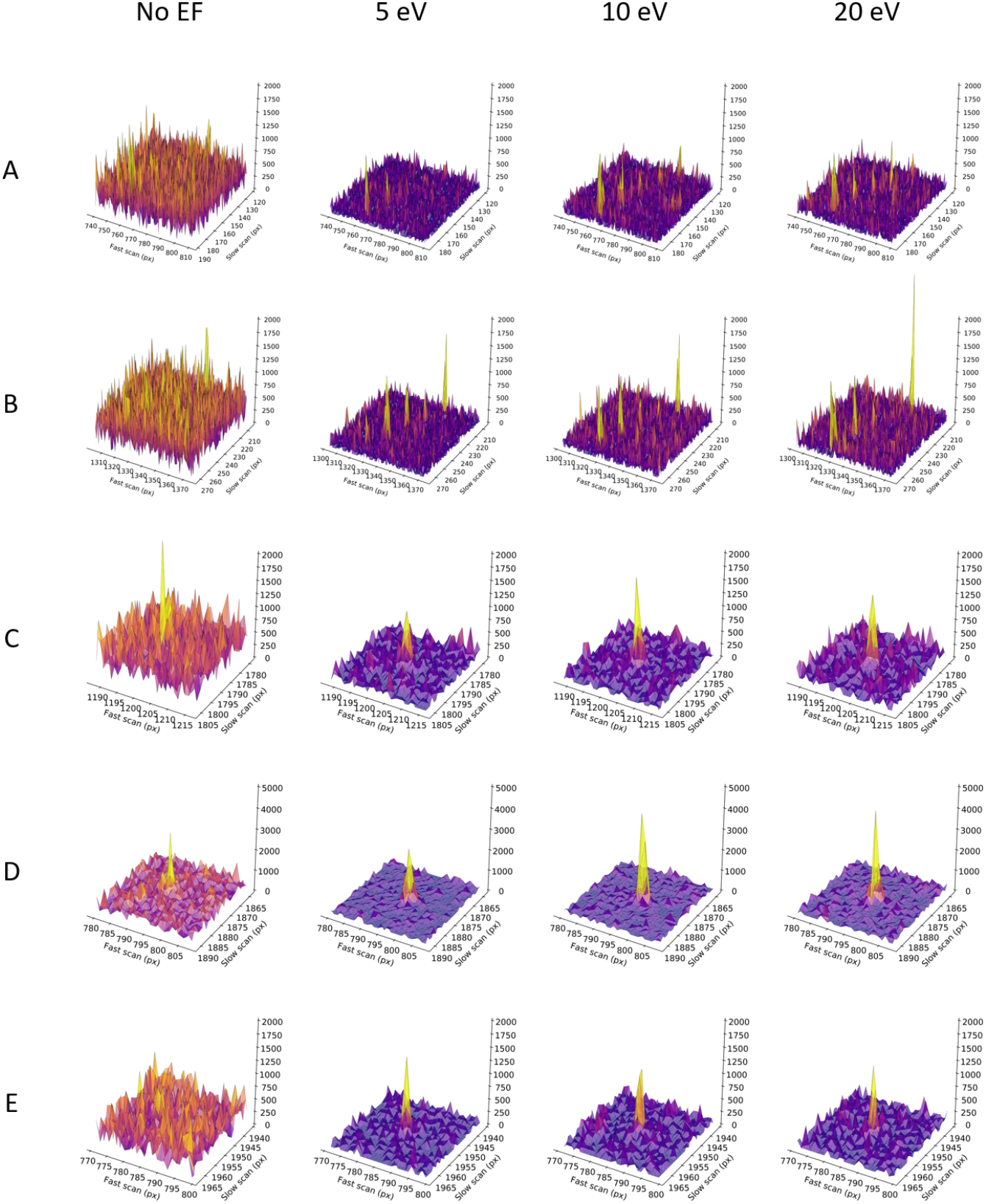
Peak profiles for unfiltered and filtered diffraction patterns taken from the same proteinase K lamella. Profiles are shown for the highlighted areas A-E in the diffraction data shown in **Figure 1.** For reference, peak profiles are shown for Bragg spots at **A**: ∼1.07 Å resolution, **B**: ∼1.17 Å resolution, **C**: 1.25 Å resolution, **D**: 1.11 Å resolution, and **E**: 1.02 Å resolution.

### 3.2. MicroED data collection at different energy filter slit widths

To further characterize the energy filter for optimal data collection, we compare intensity and model statistics for the 5, 10, and 20 eV slit widhts. Electron counting requires low flux density conditions imposed by the count-rate capability of the camera, and a low total fluence imposed by the crystal. This implies that the expected signal is proportionally weaker. Therefore, continuous rotation MicroED data were collected using an optimized strategy to maximize the signal-to-noise ratio. This approach mitigates these restrictions by rotation the crystals slower using an angular increment of 0.0476 °/s and sampling small 20.0° sweeps in reciprocal space. The total fluence for each dataset was only ∼0.84 e^-^/Å^2^, and the absorbed dose by each crystal was equivalent to ∼3.08 MGy. Nine crystals were used for MicroED data collection at each energy filter slit setting. For each slit setting, data were collected from nine lamellae to minimize variations between crystals and to ensure complete coverage of reciprocal space.

Overall, all of the filtered MicroED data show sharp and well-defined Bragg spots and low background noise. Individual datasets were processed and integrated up to a cross-correlation between two random half sets (*CC*_1/2_) that was still significant at the 0.1% level in the highest resolution shell (Karplus & Diederichs, 2012) (**Fig. 3**). All data showed strong diffraction, extending well beyond 1.2 Å resolution. Individual datasets collected with the 5 eV slit ranged between 1.18 and 1.05 Å resolution, the 10 eV data ranged from 1.13 to 1.06 Å resolution, and at 20 eV data were recorded from 1.22 to 1.06 Å resolution. Individual 10 eV datasets on average also showed higher resolution diffraction with four out of nine datasets having significant information at 1.06 Å resolution. Not all lamellae diffracted to the same resolution, which could be attributed to differences in crystallinity, quality of the lamellae, and specific lattice orientation.

**Figure 3.**
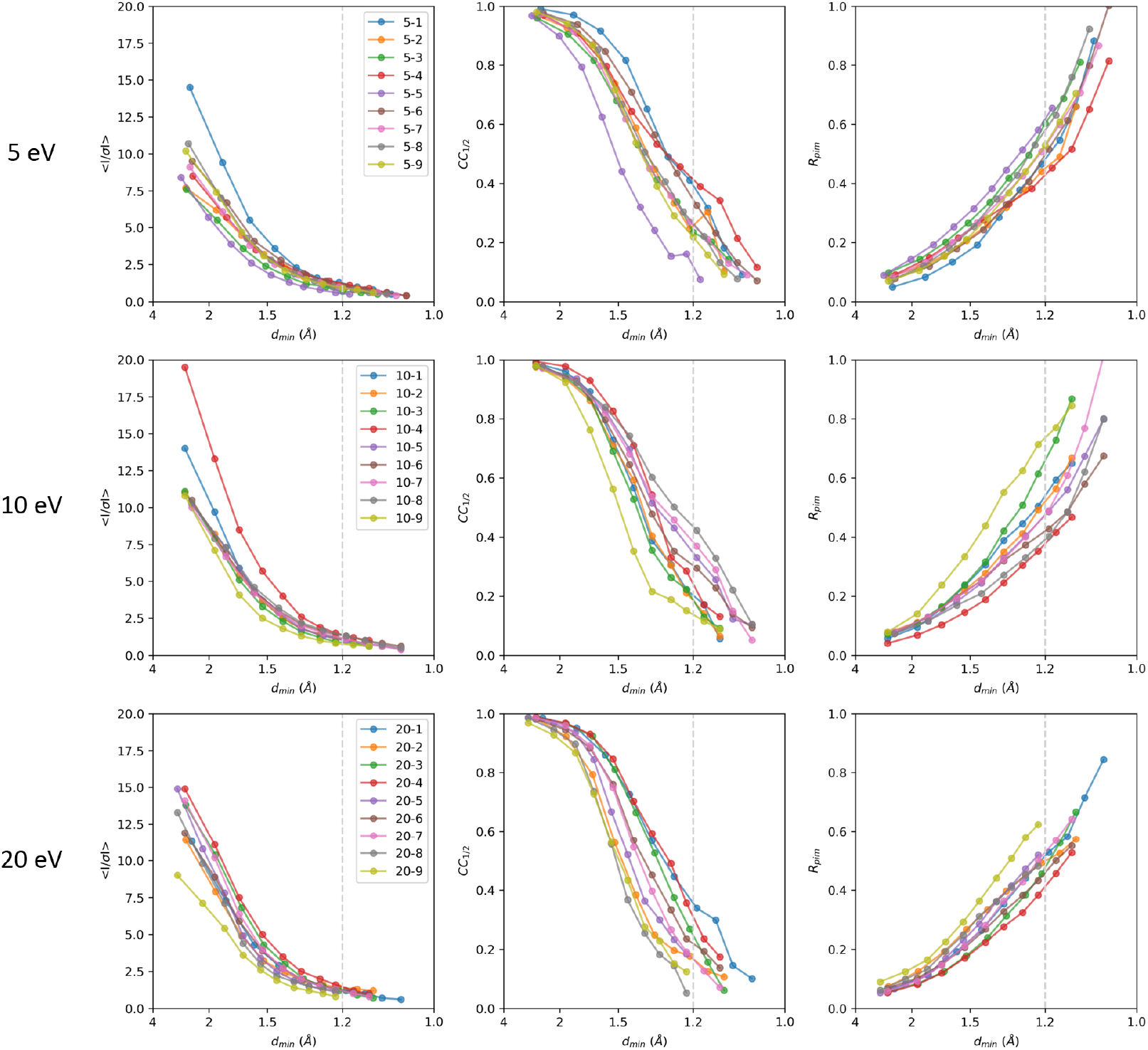
Intensity statistics for individual energy-filtered MicroED datasets collected using the 5, 10, and 20 eV slit settings. The crystallographic quality indicators mean *I/σI, CC*_1/2_, and *R*_pim_ are plotted as function of the resolution. Individual datasets were truncated at a *CC*_1/2_ that was still significant at the 0.1% level in the highest resolution shell. For a side-by-side comparison, data for each slit width were merged and truncated at 1.2 Å resolution indicated by the grey line in each plot.

Data recorded at each energy filter slit width were merged and truncated at 1.2 Å resolution, at a mean *I/σI* in the highest resolution shell of 2.0 for the 5 eV data, to enable a side-by-side comparison of the intensity and model statistics (**Table 1, Fig. 4**). In terms of signal strength, the mean *I/σI* shows a slight increase overall, as well as in the highest resolution shell, going from the narrowest 5 eV slit to the 10 and 20 eV slit settings that let electrons with a greater energy loss pass the filter (**Table 1**). Yet overall, the mean *I/σI* follows a similar trend for all slit settings, except for the lower angle reflections of the 20 eV that are stronger (**Fig. 4**). The *CC*_1/2_ for the low resolution reflections is similar between all slit settings, but then falls of rapidly for the 20 eV data to about 28.1 %, whereas the 10 eV data fall of more slowly to 44.7 %, and at 5 eV the falloff is even less to 50.5 % in the highest resolution shell (**Fig. 4**). The data have similar multiplicity and show similar values for *R*_merge_.

**Table 1.** Data merging and refinement statistics.

|  | 5 eV | 10 eV | 20 eV |
| --- | --- | --- | --- |
| **Data collection** |  |  |  |
| No. of crystals | 9 | 9 | 9 |
| Space group | *P*4<sub>3</sub>2<sub>1</sub>2 | *P*4<sub>3</sub>2<sub>1</sub>2 | *P*4<sub>3</sub>2<sub>1</sub>2 |
| Unit cell dimensions |  |  |  |
| *a*, *b*, *c* (Å) | 66.92, 66.92, 107.56 | 67.72, 67.72, 106.90 | 67.41, 67.41, 105.78 |
| $\alpha$, $\beta$, $\gamma$ (°) | 90, 90, 90 | 90, 90, 90 | 90, 90, 90 |
| Resolution (Å)<sup>†</sup> | 47.32-1.20 (1.23-1.20) | 57.21-1.20 (1.23-1.20) | 47.67-1.20 (1.23-1.20) |
| Observed reflections | 1,000,823 (76,284) | 1,262,949 (98,292) | 1,112,726 (84,484) |
| Unique reflections | 75,269 (5,504) | 76,342 (5,611) | 68,935 (5,048) |
| Multiplicity | 13.3 (13.9) | 16.5 (17.5) | 16.1 (16.7) |
| Completeness (%) | 97.9 (98.2) | 97.6 (97.9) | 89.8 (90.2) |
| *R*<sub>merge</sub> | 0.251 (0.963) | 0.262 (1.004) | 0.237 (0.936) |
| $\langle I/\sigma I \rangle$ | 6.59 (2.00) | 6.71 (2.08) | 7.18 (2.34) |
| CC<sub>1/2</sub> (%) | 99.2 (50.5) | 99.0 (44.7) | 99.1 (28.1) |
| **Refinement** |  |  |  |
| Resolution (Å) | 47.32-1.20 | 57.21-1.20 | 47.67-1.20 |
| No. of reflections | 75,213 | 76,038 | 68,168 |
| R<sub>work</sub>/R<sub>free</sub> | 0.153/0.170 | 0.167/0.182 | 0.158/0.196 |
| $\langle B \rangle$ factors (Å<sup>2</sup>) | | | |
| Overall | 10.40 | 11.13 | 10.74 |
| Protein | 8.32 | 9.05 | 8.54 |
| Ligands/ions | 9.82 | 10.54 | 8.99 |
| Waters | 24.15 | 24.87 | 25.24 |
| R.m.s. deviations |  |  |  |
| Bond lengths (Å) | 0.015 | 0.011 | 0.015 |
| Bond angles (°) | 1.21 | 1.10 | 1.24 |

<sup>†</sup>Values in parenthesis are for the highest resolution shell. Data were truncated at 1.2 Å resolution to allow for a direct comparison of intensity and model statistics between different energy filter slit settings.

**Figure 4.**
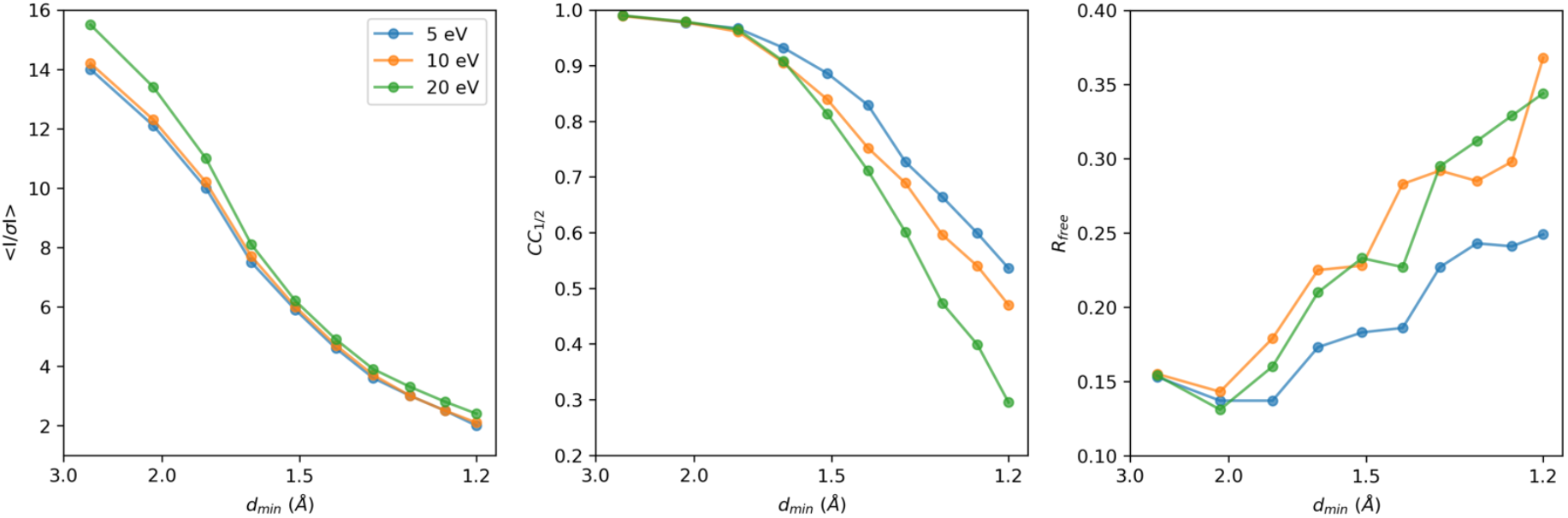
Intensity and model statistics for energy-filtered MicroED data collected using the 5, 10, and 20 eV slit widths. The mean *I/σI, CC*_1/2_, and *R*_free_ are plotted as function of the resolution. For a side-by-side comparison, data for each of the slit settings were truncated at 1.2 Å resolution.

For a direct comparison in real space, structures were refined using the same starting model and the same refinement protocol. The 5 eV data show the lowest model *R*-factors with an *R*_work_/*R*_free_ of 0.153/1.70, as well as lower *B*-factors (**Table 1**). The *R*_free_ and *CC*_free_ are better for the 5 eV data showing a less rapid falloff going towards the higher resolution shells compared that the 10 and 20 eV data that show a more similar trend (**Fig. 4**). A potential caveat is the overall completeness, which is lower for the 20 eV datasets at 89.8% (**Table 1**), which likely would not affect the intensity statistics too severely. However, the missing information could potentially affect structure refinement and map calculations. Upon visual inspection, the electrostatic potential maps did not show any noticeable differences between the different energy filter slit settings (**Fig. 5**).

**Figure 5.**
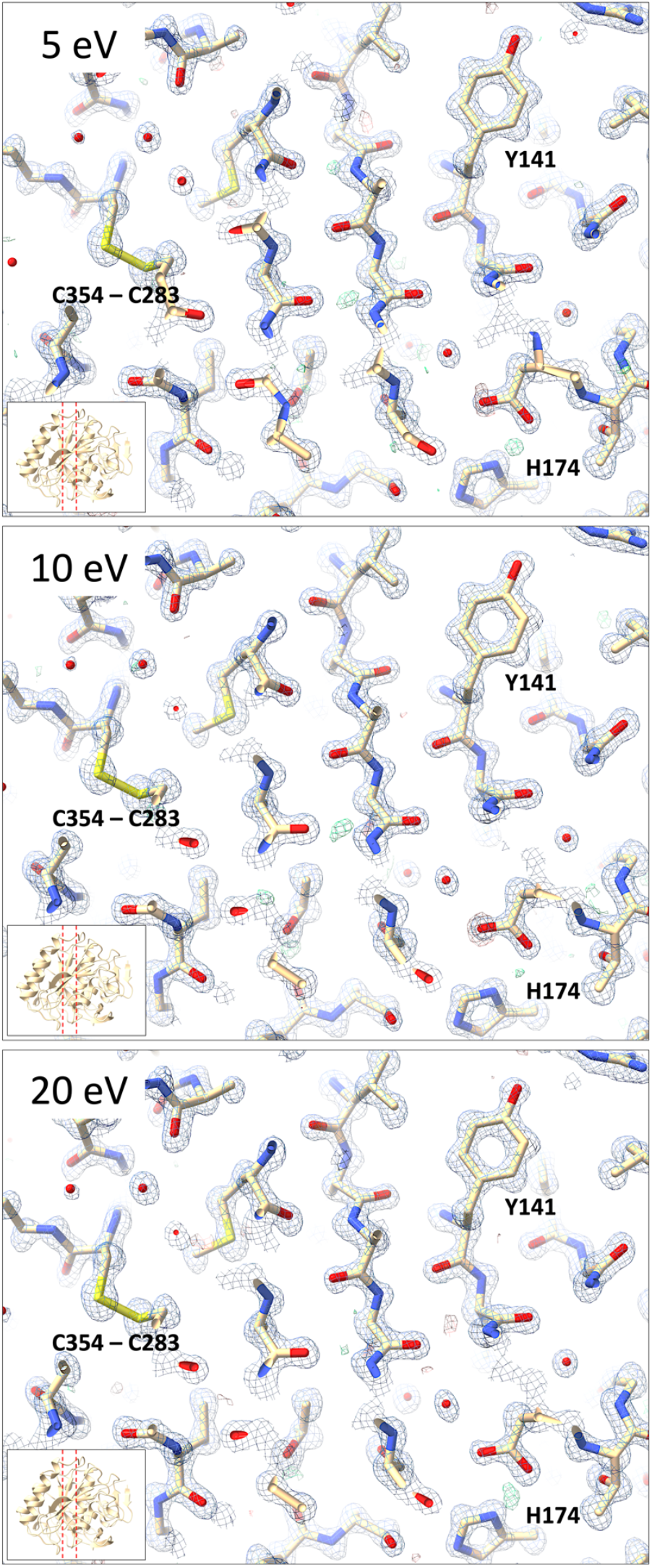
Electrostatic potential and difference maps for proteinase K from energy-filtered MicroED data using the 5, 10, and 20 eV slit settings. Data were truncated at 1.2 Å resolution. Electrostatic potential 2mFo-DFc maps are shown in blue and contoured at 2.2σ, difference mFo-DFc maps are shown in green and red for positive and negative density contoured at 3σ. No information for missing reflections was included for map calculations.

## 4. Discussion

We assess the performance of a data collection strategy combining direct electron detection with energy filtering with different slit widths to separate the inelastic noise from the elastic diffraction signal. Previously, we demonstrated a clear improvement in signal-to-noise ratio from unfiltered to energy-filtered MicroED data, leading to enhanced data quality and resolution (Clabbers *et al*., 2024). Here, we characterized the different energy filter slit widths settings by comparing diffraction stills and MicroED data (**Fig. 3**). To allow for a direct comparison, the data were all truncated at 1.2 Å resolution (**Table 1**). Even though the background noise is higher, the wider 20 eV slit settings showed the highest mean *I/σI* compared to the 5 and 10 eV data, which becomes even more pronounced for the lower resolution shells (**Fig. 4**). This can potentially be attributed to those events where electrons have undergone one or more inelastic scattering events after an initial elastic event. These electrons that are still partially coherent and focused around the Bragg spots. The wider 20 eV slit setting is less strict in separating out these partially coherent events. As the diffraction peaks become more intense, this could potentially make their detection easier as there is a stronger separation of the peaks with the background, but this might not necessarily yield more accurate intensities. In fact, the *CC*_1/2_ and R_work_/*R*_free_ are better for the 5 eV data, suggesting that the intensities that were measured with the narrower slit setting agree better internally and with the structural model (**Table 1, Fig. 4**). Additionally, there seems to be no significant loss of high-resolution information at 5 eV compared the 10 and 20 eV slit data, which on average show more intense peaks (**Fig. 3**).

These results indicate that narrower energy filter slit widths tend to perform better, suggesting there might be an advantage to exploring even narrower slit settings. Interestingly, despite showing the lowest background levels, diffraction stills and MicroED data collected using an even narrower 2 eV slit showed a significantly weaker signal compared to any of the other slit widths. On our system under the current protocol, the 2 eV slit is smaller than the SA aperture, introducing artefacts from the slit interfering with the aperture and effectively limiting the field-of-view leading to a weaker overall diffraction signal. Additionally, the narrowest 2eV slit setting potentially also cuts into part of the coherent diffraction signal as the electron source on our system has an energy spread of *ΔE* ∼ 0.8 eV. The 5 eV energy filer slit setting was the narrowest that could be used after careful alignment, causing only minimal interference at the very edges of the aperture. Selecting the wider 10 and 20 eV slits did not obscure any of the apertures, making these potential more robust to slight shifts of the energy filter slit alignment during prolonged operation. These wider slit settings show a substantial reduction in background noise compared to unfiltered data, but do not have as strong of a separation between signal and noise as observed for the 5 eV data (**Fig. 1-2**). Altogether, the 5 eV energy filter slit width provides a substantial noise reduction and retains a robust diffraction signal for reliable intensity measurements. Ultimately, the best protocol for optimal data quality differs per experiment and does not only present a trade-off between the signal and the noise, but also depends on other factors including the flux density per frame, diffraction strength of the crystals, and the radiation sensitiveness of the sample.

Our results illustrate the importance of optimizing the experimental setup and data collection strategies to allow for the best data quality. Energy filtering is an important part of enhancing data quality by selectively removing inelastically scattered electrons that contribute noise rather than a useful diffraction signal. Advances in better energy filters or more coherent electron sources can allow for even narrower slit widths, further decreasing the noise and enhancing the signal. Detector improvements enabling higher frame rates and increased dynamic range for electron counting detectors reduce acquisition time, and allow for a higher fluence per frame without sacrificing accuracy (Hattne *et al*., 2023). Additionally, data collection strategies that combine even more small sweeps (Yonekura *et al*., 2015), or adopting a serial crystallography approach that uses snapshots of many individual crystals (Bücker *et al*., 2020), could provide a significant boost in attainable resolution. These approaches could increase redundancy and improve the accuracy of intensity measurements by better sampling reciprocal space, thus minimizing errors due to mechanical drift or crystal misalignment over longer data collection periods. By systematically refining data collection parameters such as energy filter slit width, fluence per frame, and total accumulated dose, MicroED experiments can be optimized for specific sample types, enhancing signal-to-noise ratio and resolution in a controlled and reproducible way. Ultimately, optimal adjustment of the data collection parameters leads to better data quality, which informs better structural models.

## Data availability

Coordinates and structure factors have been deposited in the Protein Data Bank (PDB) under accession codes XXXX [https://doi.org/10.2210/pdbXXXX/pdb] (Proteinase K, 5 eV energy filter slit), XXXX [https://doi.org/10.2210/pdbXXXX/pdb] (Proteinase K, 10 eV energy filter slit), XXXX [https://doi.org/10.2210/pdbXXXX/pdb] (Proteinase K, 20 eV energy filter slit). EM maps have been deposited in the Electron Microscopy Data Bank (EMDB) under accession codes EMD-XXXXX [https://www.ebi.ac.uk/pdbe/entry/emdb/EMD-XXXXX] (Proteinase K, 5 eV energy filter slit), EMD-XXXXX [https://www.ebi.ac.uk/pdbe/entry/emdb/EMD-XXXXX] (Proteinase K, 10 eV energy filter slit), EMD-XXXXX [https://www.ebi.ac.uk/pdbe/entry/emdb/EMD-XXXXX] (Proteinase K, 20 eV energy filter slit).

## Acknowledgements

This study was supported by the National Institutes of Health (P41GM136508), the Department of Defense (HDTRA1-21-1-0004), and the Howard Hughes Medical Institute. Portions of this research or manuscript completion were developed with funding from the Department of Defense MCDC-2202-002. Effort sponsored by the U.S. Government under Other Transaction number W15QKN-16-9-1002 between the MCDC, and the Government. The US Government is authorized to reproduce and distribute reprints for Governmental purposes, notwithstanding any copyright notation thereon. The views and conclusions contained herein are those of the authors and should not be interpreted as necessarily representing the official policies or endorsements, either expressed or implied, of the U.S. Government. The PAH shall flow down these requirements to its sub awardees, at all tiers.

